# T_2_ relaxometry for myelin water fraction: an ex vivo brain imaging study

**DOI:** 10.64898/2026.09.14.751605

**Authors:** Liana Sanches, Narges Taghizadehsalehabad, Roqaie Moqadam, Walter Adame-Gonzalez, Zaki Alasmar, Dominique Mirault, Gian Franco Piredda, Gustavo Turecki, Josefina Maranzano, Naguib Mechawar, The CIMA-Q group, Mallar Chakravarty, Mahsa Dadar, Yashar Zeighami

**Author notes:** Corresponding Authors: Liana Sanches, Cerebral Imaging Centre, Douglas Research Centre, 6875 Boulevard LaSalle, Montréal, QC, H4H 1R3., Mahsa Dadar, Cerebral Imaging Centre, Douglas Research Centre, 6875 Boulevard LaSalle, Montréal, QC, H4H 1R3., Yashar Zeighami, Cerebral Imaging Centre, Douglas Research Centre, 6875 Boulevard LaSalle, Montréal, QC, H4H 1R3. Co-first authors (Liana Sanches, Narges Taghizadehsalehabad). Co-senior authors (Mahsa Dadar, Yashar Zeighami). Data used in the preparation of this article were obtained from the Consortium for the early identification of Alzheimer’s disease – Quebec (CIMA-Q; cima-q.ca). A list of researchers involved in the design of CIMA-Q can be found on the cima-q.ca website. These researchers contributed to the establishment of protocols, the implementation of the research infrastructure, the recruitment and follow-up of participants, the obtaining of data, the maintenance of biological and ex-vivo samples, and certain derived data.

## Abstract

**Introduction:** Myelin water imaging provides a non-invasive approach for indirectly investigating myelin content of the brain by magnetic resonance Imaging (MRI). Myelin water fraction (MWF) reflects the fraction of water signal associated with water trapped between myelin layers. Postmortem imaging provides a unique opportunity to validate MWF metrics against gold-standard postmortem neuropathology. However, there is a need for validated sequences that would be applicable in postmortem settings. In this study, we validated and compared the applicability of turbo spin echo (TSE) and gradient-and-spin-echo (GRASE) techniques for myelin water imaging in formaldehyde fixed postmortem human brains.

**Methods:** 40 postmortem human brain hemispheres were scanned with TSE and GRASE sequences. Acquired data were reconstructed using a non-parametric multicomponent T_2_ relaxometry approach using a consistent framework to support comparison between GRASE- and TSE-derived measures. Agreement between derived measures was quantified at both voxel and regional level. Maps derived from the full 32-echo GRASE reconstruction were compared against a truncated 14-echo reconstruction to examine the contribution of the longer echoes to the derived maps. Finally, a random forest regression model was trained based on TSE data to predict the GRASE-derived MWF maps.

**Results:** Both sequences provided robust multicomponent T_2_ characterization across the brain, and the derived measures showed anatomical patterns consistent with expected differences across tissue types. TSE and GRASE derived maps had moderate to strong agreement at voxel (0.41< ρ < 0.82, all p_FDR_<0.001) and regional levels (0.87< ρ < 0.98, all p_FDR_<0.001), with consistently stronger agreement observed for regional metrics. Echo reduction comparisons between the full 32-echo GRASE reconstruction and a truncated 14-echo reconstruction showed minimal impact of echo truncation (all ρ > 0.98, p_FDR_ < 0.001). The nonlinear random forest regression model trained on TSE maps was able to accurately reproduce the MWF maps derived from GRASE (r = 0.91, RMSE = 8.87).

**Discussion:** TSE and GRASE32 capture strongly related multicomponent T₂ information in fixed postmortem human brain tissue, but with systematic and tissue-dependent quantitative differences across derived measures. These differences suggest that direct interchangeability should not be assumed and that cross-sequence mapping or calibration is required when quantitative equivalence is desired.

## Introduction

Magnetic resonance imaging (MRI) relaxometry can be used to investigate myelin integrity by characterizing the relaxation properties of water associated with myelin. More specifically, T_2_ relaxation times reflect the local microenvironment and tissue composition surrounding water protons and are sensitive to tissue water content (1). In vivo, the short-T_2_ component in healthy white matter (WM) is typically observed at approximately 10–40 ms and is commonly attributed to water trapped between the layers of the myelin sheath (2). The myelin water fraction (MWF) is defined as the signal fraction arising from this short-T_2_ component relative to the total water signal (2). An intermediate-T_2_ component, typically characterized by relaxation times of approximately 40–200 ms, is generally associated with intra- and extra-axonal/extramyelinic water. Although less extensively studied than MWF, alterations in the intermediate-T_2_ component have been associated with changes in tissue water content during aging (3) as well as pathological processes in individuals with subjective cognitive decline (4). The longest-T_2_ component, with relaxation times typically on the order of seconds, is generally attributed to freely mobile water, including the cerebrospinal fluid (CSF) (5).

Quantifying these components requires sampling the T_2_ decay over multiple echo times. Conventional multi-echo spin-echo (MESE)(6) have formed the basis of multicomponent T_2_ imaging and modeling. However, acquiring a sufficiently long and densely sampled echo train, while maintaining adequate signal-to-noise ratio (SNR), spatial resolution, and brain coverage can result in long acquisition times, limiting feasibility in both research and clinical settings and increasing sensitivity to subject motion. In addition, factors inherent to MESE acquisitions, including imperfect refocusing across the echo train, can generate stimulated echoes and other non-ideal signal pathways (7). These effects alter the measured echo amplitudes and complicate quantitative T_2_ estimation. These limitations have motivated the development of faster acquisition strategies for myelin water imaging.

3D gradient-and-spin-echo (GRASE) imaging was introduced as one such approach. The sequence developed by Prasloski et al. (8) enabled whole-cerebrum myelin water imaging in less than 15 minutes, substantially improving its feasibility compared with conventional MESE approaches (8). More recently, Piredda et al. (9) proposed a further accelerated 3D multi-echo GRASE using CAIPIRINHA parallel imaging, achieving whole-brain MWF mapping at 1.6-mm isotropic resolution in approximately 8 minutes. The use of GRASE sequences for MWF estimation has been previously investigated and validated against established 2D (8) and 3D MESE approaches (7,9). However, at this time, the GRASE sequence is not freely available and may require specific research agreements, which can limit its broader use. A potential alternative is to use the accelerated version of the spin-echo sequence, called turbo spin echo (TSE). A train of 180° refocusing pulses generates multiple spin echoes within a single TR, allowing substantially reducing acquisition time. Separated single-echo sequences reduce the problem of stimulated echoes while flip angle and B1 effects can be incorporated to the modelling using Extended Phase Graphic (EPG) -based signal models (10). Such an approach has the advantage of using a widely available sequence while still providing multiple samples of the T_2_ decay.

On the analytical side, multicomponent T_2_ models represent the observed signal decay as a combination of water pools with distinct T_2_ relaxation times and are commonly analyzed using multiexponential fitting approaches (11). This allows the measured signal to be characterized in terms of distinct tissue water compartments, providing a framework for estimating their relative contributions. Because the estimation of multicomponent T_2_ distributions is an ill-posed inverse problem, the recovered distribution can depend on both the reconstruction approach and characteristics of the acquired data (1,12,13). Consequently, the recovered T_2_ distribution may be influenced by acquisition characteristics such as echo sampling, SNR, and sequence-specific signal formation, highlighting the importance of direct comparison of these measures across acquisition strategies.

Postmortem brain tissue provides an invaluable setting for cross-sequence comparisons, where the same specimen can be scanned using multiple acquisition protocols under controlled conditions, free of motion and with longer scan-durations. Furthermore, for ex vivo relaxometry, compared to T1, T_2_ measurements are less strongly influenced by formalin fixation and tissue depth, which favors a postmortem acquisition comparison for T_2_ relaxometry. However, formalin fixation still alters T_2_ relaxation properties and therefore remains an important consideration. In five human brain specimens, Dawe et al. reported that fixation-related changes in T_2_ are spatially heterogeneous and time dependent, likely reflecting progressive formaldehyde penetration and subsequent tissue changes during prolonged fixation (14). Using 3D GRASE, Shatil et al. (15) reported a progressive pattern of formalin fixation in two human brain specimens, with both T1 and T_2_ decreasing with increasing fixation time, while MWF increased, consistent with a fixation-related shift toward shorter T_2_ components. In our prior work using a substantially larger postmortem cohort, we similarly found that progressive formaldehyde fixation produced systematic changes in relaxation-related MRI measures, including decreasing T_2_^∗^ and increasing MWF, with distinct spatial and temporal trajectories across the brain (16). Together, these findings indicate that T_2_ relaxometry is comparatively well suited for cross-sequence evaluation in postmortem tissue; nevertheless, fixation duration remains an important source of variability that should be considered when comparing acquisition strategies.

Establishing a feasible and widely available acquisition for myelin water imaging in postmortem tissue raises several questions. First, given the differences between in vivo and ex vivo tissue, particularly the shortening of T₂ with fixation, it has not been systematically established whether the extended echo coverage commonly used for in vivo multicomponent T₂ imaging is necessary in fixed specimens. It therefore remains unclear whether a shorter echo train can preserve the principal myelin-related measures obtained from a standard 32-echo GRASE acquisition. Second, although multi-echo TSE provides a more widely available approach for densely sampling T_2_ decay, it has not been established whether ME-TSE can recover multicomponent T_2_ information comparable to that obtained with GRASE in fixed human brain tissue, or whether the relationship between the two sequences varies across tissue types. Given the effects of fixation on T_2_ relaxation and MWF, it is also unclear whether fixation-related changes are captured similarly by the two sequences. Where the two sequences differ, it is unclear whether these differences reflect a loss of myelin-related information or sequence-dependent differences in how the recovered T_2_ distribution is separated and partitioned into components, and whether the relationship between the two acquisitions can be modeled.

In the present study, we directly compared 3D GRASE and ME-TSE in formaldehyde-fixed postmortem human brain hemispheres across a broad range of fixation durations. We assessed whether reducing the 32-echo GRASE acquisition to a 14-echo range comparable to that sampled by ME-TSE preserved the principal multicomponent T_2_ measures, including MWF, IEWF, T_2,M_, and T_2,IE_. We also directly compared the 15-echo ME-TSE and GRASE32 acquisitions in the same specimens at voxelwise, regional, and tissue-class levels to characterize the magnitude and tissue dependence of cross-sequence differences. We further examined the effects of fixation duration on T_2_ distributions and MWF and whether these effects differed between the two sequences. Finally, we investigated whether the relationship between the two acquisitions could be modeled and to what degree the GRASE32 MWF could be predicted from TSE-derived multicomponent measures and from the full TSE T_2_ distribution. Together, these analyses aimed to determine whether reduced echo sampling is sufficient for multicomponent T_2_ characterization in fixed postmortem tissue and whether a more widely available ME-TSE acquisition can capture the myelin-related information provided by GRASE.

## Methods

Forty postmortem human brain hemispheres obtained from the Douglas Brain Bank (DBB; https://douglasbrainbank.ca/) were used in this study. The sample included specimens without neurological disorders as well as specimens from donors with different neurodegenerative diseases including Alzheimer’s disease (AD), Parkinson’s disease (PD), frontotemporal dementia (FTD), vascular dementia (VD), Lewy body dementia (LBD), and amyotrophic lateral sclerosis (ALS). Brain specimens are collected in accordance with informed consent from the donors or their next of kin, according to tissue banking practices regulated by the Quebec Health Research Fund and the Guidelines on Human Biobanks and Genetic Research Databases overseen by the Douglas Research Ethics Board (17). Ethics approval for the study was obtained from the Ethics Review Board of the Douglas Research Centre.

Tissue processing and fixation procedures have been previously described (17). Following processing, intact hemispheres are immersed in 10% formalin in airtight MRI-compatible plastic containers for long-term preservation. The included specimen spanned a wide range of formaldehyde fixation durations to allow for characterization of the impact of fixation on derived metrics (16). A subset of the specimens (N = 10) were from participants in the Consortium for the Early Identification of Alzheimer’s Disease-Quebec (CIMA-Q) (18).

## MRI acquisition

MRI acquisitions were performed on a 3T Siemens Prisma Fit scanner at the Douglas Cerebral Imaging Centre. Specimen containers were positioned within a 64-channel head/neck coil. Within each container, the hemisphere was positioned flat on the sagittal cut, with the cerebellum oriented toward the bore, and stabilized in place using a custom-designed 3D-printed grid (16) to standardize positioning and minimize motion-related artifacts. Structural T1-weighted (T1w) and T2-SPACE images were acquired for anatomical reference and tissue segmentation (19). Multi-echo T_2_-weighted imaging included a 2D multi-echo TSE sequence and a 3D GRASE sequence (Table 1).

**Table 1.**
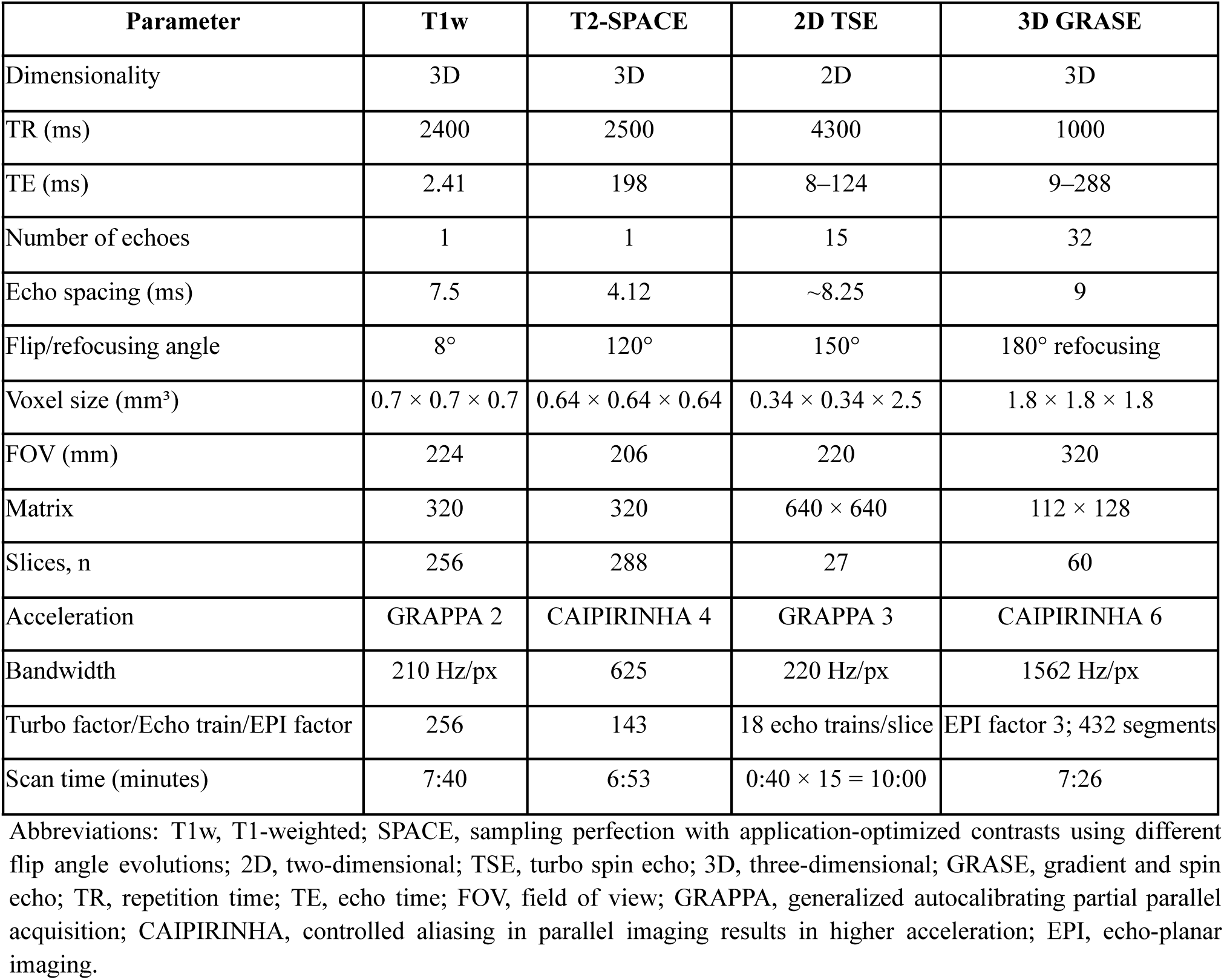
MRI acquisition protocol.

## Image Preprocessing and Tissue Segmentation

T1w and T2-SPACE structural images were preprocessed and linearly registered to the stereotaxic space (17). SynthSeg (20) was applied to the processed T1w images to generate initial tissue labels, and left and right hemisphere labels were merged. Since SynthSeg does not label white matter hyperintensities (WMH), these were manually segmented and added to the existing labels. The Hammers lobar atlas was used to derive lobar cortical gray matter (cGM) and white matter (WM) segmentations. The final regional label set included lobar cGM and WM regions, manually defined lobar WMHs, cerebellar GM and WM, and subcortical GM and brainstem structures. Regional labels were then used to summarize the quantitative measures within each anatomical region.

## Multicomponent T_2_ reconstruction

Multi-echo T_2_-weighted data were reconstructed using a non-parametric multicomponent T_2_ relaxometry approach. T_2_ distributions were estimated using the Python multicomponent-T2-toolbox (13) (https://github.com/ejcanalesr/multicomponent-T2-toolbox/), which provides several approaches for estimating T_2_ spectra, including unregularized non-negative least squares (NNLS) and multiple regularized NNLS methods based on different regularization criteria and penalty matrices. We used Bayesian regularized non-negative least squares (BayesReg) with an L2 regularization matrix.

For each voxel, the fitted T_2_ distribution was used to derive quantitative maps, including MWF, intra-/extra-cellular water fraction (IEWF), free-water fraction (FWF). MWF, IEWF, and FWF were defined as the relative spectral amplitudes within predefined short-, intermediate-, and long-T_2_ intervals, respectively (21). The geometric mean T_2_ of the myelin-water component (T_2,M_), geometric mean T_2_ of the intra-/extra-cellular component (T_2,IE_) were also calculated. The same reconstruction framework and component definitions were applied across sequences to support direct comparison between GRASE- and TSE-derived measures.

Implementation parameters were kept the same across reconstruction variants. Reconstructions were performed without tissue masking. GRASE reconstructions used a minimum echo time of 9 ms and TR of 1000 ms, whereas TSE reconstructions used a minimum echo time of 8 ms and TR of 4300 ms. Effective refocusing flip-angle estimation was performed using the spline method with flip-angle smoothing enabled. T_2_ distributions were estimated using a logarithmically spaced T_2_ dictionary ranging from 8 to 2000 ms with 60 components. The component intervals were defined as 10 ≤ *T* _2_< 40 ms for MWF, 40 ≤ *T* _2_< 200 ms for IEWF, and *T* _2_≥ 200 ms for FWF. Fraction maps, including MWF, IEWF, and FWF, were multiplied by 100 to express values as percentages, whereas T_2,M_, T_2,IE_ and other T_2_-derived measures were retained in milliseconds. The final maps were used for spatial resampling, voxel-wise agreement analyses, and regional quantification.

## Sequence Comparisons Echo-subset reconstruction

Multicomponent T_2_ measures were compared between the full GRASE and TSE acquisitions, with GRASE32 versus TSE serving as the primary sequence comparison. Because GRASE32 sampled a broader TE range than TSE, differences between the two protocols could reflect both sequence-related effects and differences in echo coverage. To evaluate the contribution of echo coverage, an additional reduced GRASE reconstruction was generated from the early echoes of the full GRASE acquisition. GRASE14 was reconstructed using the first 14 echoes, corresponding to a TE range of 9–126 ms, which closely matched the TE range of TSE15 (8–124 ms), providing a within-sequence assessment of the effect of echo truncation (Table 2). For the primary analysis, GRASE32 was compared with GRASE14 and TSE to assess agreement across sequences. These comparisons were carried out both voxel-wise and at the level of predefined anatomical regions.

**Table 2.** Reconstruction variants used for echo-subset and sequence-comparison analyses.

| Reconstruction | Sequence | N Echoe | TE range (ms) | Details |
| --- | --- | --- | --- | --- |
| GRASE32 | 3D GRASE | 32 | 9–288 | Full GRASE acquisition |
| GRASE14 | 3D GRASE | 14 | 9–126 | Early-echo subset (TE range matching TSE) |
| TSE | 2D multi-echo TSE | 15 | 8–124 | Full TSE acquisition |

## Voxelwise and Regional Comparisons

Voxel-wise analyses required the quantitative maps to occupy the same common space. GRASE32 and GRASE14 shared the same acquisition grid and required no resampling. TSE-derived maps, however, differed from GRASE in voxel size, orientation, and image geometry. The higher-resolution TSE maps were therefore resampled onto the GRASE grid (itk_resample, first order polynomial option), using GRASE geometry as the reference (22,23). Anatomical label maps were also resampled to the native GRASE and TSE spaces (itk_resample, label resampling option using nearest neighbor interpolation). Regional values for GRASE32, GRASE14, and TSE were then extracted using their corresponding label maps. For each specimen, sequence, and reconstruction variant, mean values were extracted within each region for the derived quantitative maps (MWF, IEWF, T_2,M_, and T_2,IE_).

## Statistical Analyses

For each quantitative measure, agreement between reconstruction variants (i.e., GRASE vs echo truncated GRASE) and acquisition sequences (i.e., GRASE vs TSE) was quantified using Pearson correlation, Spearman rank correlation, mean difference, mean absolute error, and root mean squared error, computed within binary brain masks for voxel-wise comparisons and within each anatomical region for regional comparisons. For voxel-wise comparisons, linear regression was additionally used to characterize the relationship between paired measurements. Analyses were performed separately for each derived measure, including MWF, IEWF, T_2,M_, and T_2,IE_.

To determine whether GRASE32–TSE associations differed by tissue class, separate linear mixed-effects models were fitted for each derived measure, with GRASE32 as the dependent variable and TSE, tissue class, and their interaction as fixed effects, with specimen ID included as a random intercept. The contribution of the TSE × tissue interaction was assessed by comparing model fit with and without the interaction term, using a likelihood-ratio test. Tissue-specific slopes were derived from the fixed effects of the full interaction model. Spearman rank correlations were also calculated separately within each tissue class to assess the strength of the association between TSE- and GRASE32-derived measurements. For analyses involving multiple comparisons, p-values were adjusted using the Benjamini–Hochberg false discovery rate (FDR) procedure.

## Signal-background quality assessment

Signal-to-noise ratio (SNR) was evaluated independently for each sequence, since differences in acquisition geometry, parallel imaging, reconstruction, and receiver-coil combination confounds direct comparison of absolute SNR (23) between TSE and GRASE. The goal was to confirm whether each acquisition maintained adequate signal relative to its own noise levels across the sampled echo times. For each specimen and sequence, measurements were extracted across echo times from tissue, formalin, and air regions. Tissue signal was measured within a binary tissue mask generated from the anatomical label map in the corresponding sequence space. Formalin background was sampled using five manually placed ROIs per specimen. Air background was sampled using ROIs outside the container. All ROIs were visually inspected to confirm that they were confined to the intended tissue compartment and did not include adjacent tissue types.

SNR measures were calculated as the mean tissue signal divided by the standard deviation of the background region for each echo. To characterize signal decay within each sequence, tissue signal was normalized within each specimen and sequence by the signal at the first echo time:

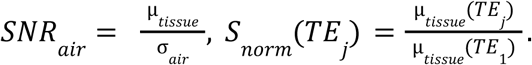

### Impact of long-term formaldehyde exposure on T_2_ distributions and MWF

To characterize the effect of prolonged formaldehyde exposure on multicomponent T_2_ estimates, whole-brain T_2_ distributions were examined separately for GRASE32 and TSE. For each specimen, the regional mean T_2_ spectrum was normalized to its total spectral amplitude and expressed as a percentage of the full T_2_ distribution. The 40-ms cutoff threshold used to define the short-T2 component was indicated on each distribution. The association between fixation duration and whole-brain MWF was assessed using a linear mixed-effects model. Fixation duration was first transformed as log_10_(fixation days+1) to account for its wide temporal range and then standardized across specimens. MWF values from both sequences were standardized using a common pooled mean and standard deviation. The model included fixation duration, acquisition sequence, and their interaction as fixed effects, with age and sex included as covariates and a specimen-specific random intercept to account for paired GRASE32 and TSE measurements. GRASE32 was used as the reference sequence. Sequence-specific standardized fixation slopes were obtained from the model, and the fixation-by-sequence interaction was used to test whether the association between fixation duration and MWF differed between GRASE32 and TSE.

## Cross-sequence MWF prediction

To evaluate whether multicomponent T_2_ information derived from the TSE acquisition could be used to predict GRASE-derived MWF, we performed voxel-wise cross-sequence prediction using linear and random forest regression models (24). Predictors were derived from the multicomponent T_2_ analysis of the TSE data to predict the voxelwise GRASE32 MWF maps. To establish voxel-wise correspondence between acquisitions, TSE-derived maps and T_2_ distributions were resampled to the GRASE32 image space prior to modeling as previously described. Regional anatomical information was included in all models as an independent feature to account for tissue-and region-dependent differences in the relationship between TSE- and GRASE-derived MWF.

Initially, two linear regression models were evaluated (Table 3). The first modelled GRASE32 as a linear function of TSE-derived MWF with regional anatomy included as a categorical predictor. The second additionally included a quadratic TSE MWF term to capture nonlinear cross-sequence relationships. Four random forest models were evaluated to determine which TSE-derived information combination resulted in the best prediction for GRASE-derived MWF; Regional anatomy information was included as a categorical predictor in all models. In addition to the regional anatomy, Model 1 included TSE-derived MWF alone. Model 2 incorporated the four T2-derived measures (MWF, IEWF, T_2,M_, and T_2,IE_). Model 3 used principal components (25) derived from the normalized TSE T_2_ distribution; and Model 4 combined the spectral principal components with TSE-derived MWF and T_2,M_. The number of principal components was determined through sensitivity analysis, with N = 12 selected for the final spectrum-based models. Random forest regression was implemented using 100 trees, a minimum leaf size of 500 voxels, three predictors sampled at each split, and an in-bag fraction of 0.70.

**Table 3.** Predictor configurations used for cross-sequence prediction of GRASE32-derived MWF from TSE-derived information.

| Model | Predictor set |
| --- | --- |
| **Linear Model** | $MWF_{\text{GRASE}} \sim MWF_{\text{TSE}} + R$ |
| **Quadratic Model** | $MWF_{\text{GRASE}} \sim MWF_{\text{TSE}} + (MWF_{\text{TSE}})^2 + R$ |
| **RF1: MWF** | $MWF_{\text{GRASE}} \sim \text{RF}(MWF_{\text{TSE}}, R)$ |
| **RF2: Four derived measures** | $MWF_{\text{GRASE}} \sim \text{RF}(MWF_{\text{TSE}}, IEWF_{\text{TSE}}, T_{2,M, \text{TSE}}, T_{2,IE, \text{TSE}}, R)$ |
| **RF3: Spectrum** | $MWF_{\text{GRASE}} \sim \text{RF}(PC1:12; R)$ |
| **RF4: Combined** | $MWF_{\text{GRASE}} \sim \text{RF}(PC1:12, MWF_{\text{TSE}}, T_{2,M, \text{TSE}}, R)$ |

Abbreviations: RF, random forest; R, region (categorical predictor); PC1–12, first 12 principal components of the normalized TSE $T_2$ distribution.

Model performance was evaluated using 10-fold specimen-level cross-validation (26), such that all voxels from a given specimen were assigned exclusively to either the training or held-out set within each fold. The same fold assignments were used across all model configurations to enable direct comparison of predictive performance. Random forest models were trained using specimen-balanced voxel weights so that specimens contributing larger numbers of within mask voxels did not disproportionately influence model fitting. Predictions for each specimen were generated only from the model for which that specimen was held out. Linear regression models were evaluated using the same specimen-wise cross-validation folds as the random forest models.

For models incorporating the T_2_ distribution as a predictor, the TSE spectrum was first spatially resampled to the GRASE32 space, with each spectral bin resampled independently using linear interpolation. The resampled spectrum was then normalized voxel-wise by its total spectral amplitude to represent the relative distribution of T_2_ components independent of the overall signal magnitude. Principal component analysis (PCA) was used to reduce the dimensionality of the normalized spectra. To prevent information leakage, PCA was applied to only the training specimens within each cross-validation fold, and the resulting transformation was projected to the corresponding held-out specimens to derive the corresponding features and prediction.

Model performance was quantified separately for each held-out specimen by comparing predicted and observed GRASE32 MWF values on the common set of valid voxels. Prediction error was summarized using root mean squared error (RMSE) and mean absolute error (MAE) (27), while agreement was assessed using the concordance correlation coefficient (CCC) (28), mean prediction bias, and the ratio of the predicted to observed standard deviation as a measure of dynamic-range preservation. Spatial similarity was additionally assessed using masked structural similarity index (SSIM) (29). Model performance was summarized across specimens using the median of each metric.

## Results

Table 4 summarizes the characteristics of the included specimens. 40 specimens were acquired with complete GRASE32 and TSE15 acquisitions and were included in the sequence-comparison analyses. The analyzed specimens had a mean age at death of 79.8 ± 11.3 years, and 17 (42%) were female. Mean PMI was 33.7 ± 21.3 hours, and mean fixation duration was 9.6 ± 7.0 years. The sample included a range of neurodegenerative diagnoses as well as specimens without neurological disorders (Table 4). Representative GRASE32 and TSE15 maps are shown in Figure 1.

**Figure 1.**
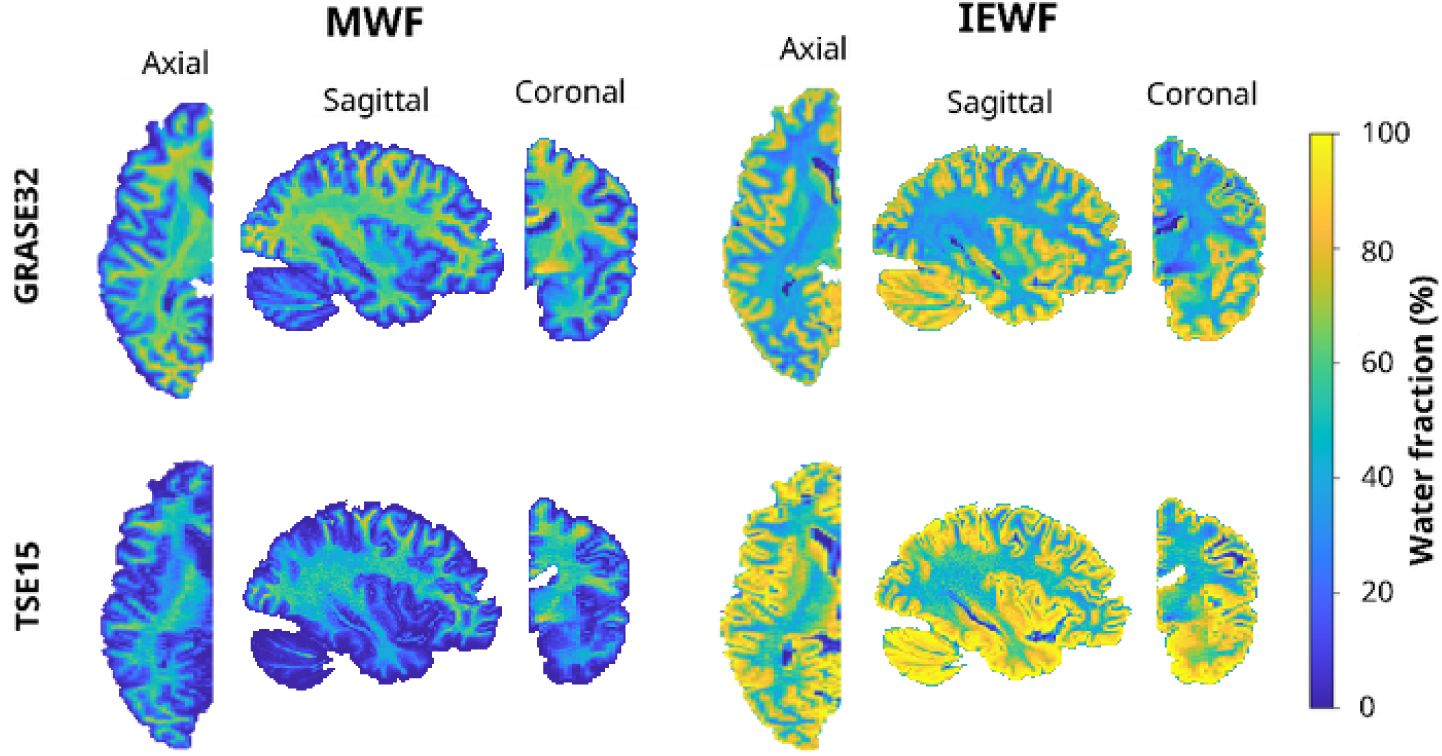
Representative myelin water fraction (MWF) and intra-/extracellular water fraction (IEWF) maps obtained with GRASE32 and TSE15. Sagittal, coronal, and axial views are shown for MWF (left) and IEWF (right), with GRASE32 in the top row and TSE15 in the bottom row. Maps were estimated using BayesReg-L2 with a 40-ms myelin-water cutoff and a minimum T_2_ of 8 ms, and are displayed in their native acquisition spaces without anatomical masking. A common color scale of 0–100% is used for both water-fraction measures to facilitate visual comparison between acquisitions.

**Table 4.**
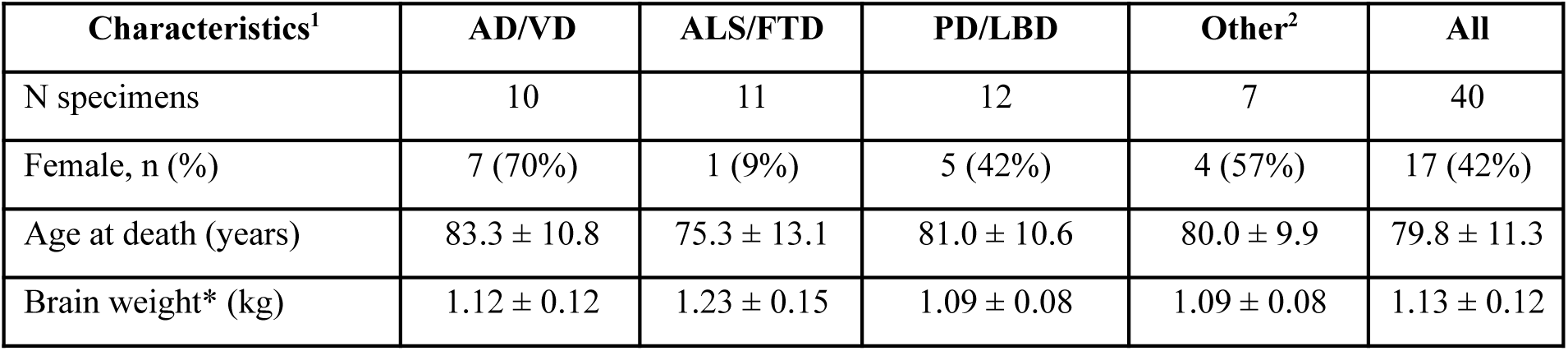
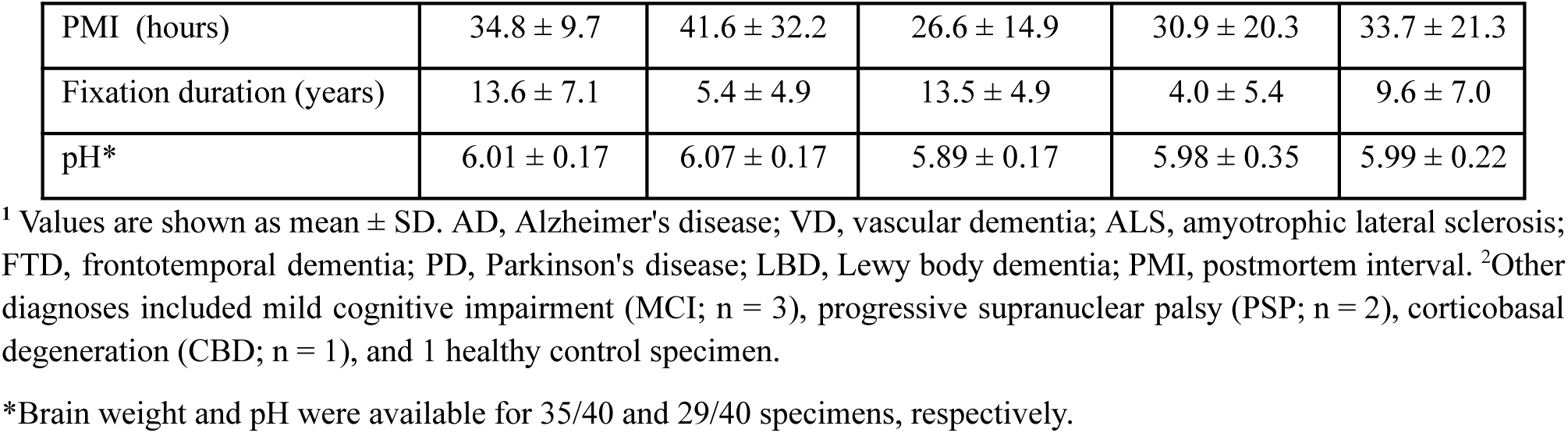
Demographic, diagnostic, and postmortem characteristics of included specimens.

| Characteristics<sup>1</sup> | AD/VD | ALS/FTD | PD/LBD | Other<sup>2</sup> | All |
| --- | --- | --- | --- | --- | --- |
| N specimens | 10 | 11 | 12 | 7 | 40 |
| Female, n (%) | 7 (70%) | 1 (9%) | 5 (42%) | 4 (57%) | 17 (42%) |
| Age at death (years) | $83.3 \pm 10.8$ | $75.3 \pm 13.1$ | $81.0 \pm 10.6$ | $80.0 \pm 9.9$ | $79.8 \pm 11.3$ |
| Brain weight* (kg) | $1.12 \pm 0.12$ | $1.23 \pm 0.15$ | $1.09 \pm 0.08$ | $1.09 \pm 0.08$ | $1.13 \pm 0.12$ |
| PMI (hours) | 34.8 ± 9.7 | 41.6 ± 32.2 | 26.6 ± 14.9 | 30.9 ± 20.3 | 33.7 ± 21.3 |
| Fixation duration (years) | 13.6 ± 7.1 | 5.4 ± 4.9 | 13.5 ± 4.9 | 4.0 ± 5.4 | 9.6 ± 7.0 |
| pH* | 6.01 ± 0.17 | 6.07 ± 0.17 | 5.89 ± 0.17 | 5.98 ± 0.35 | 5.99 ± 0.22 |

<sup>1</sup> Values are shown as mean ± SD. AD, Alzheimer's disease; VD, vascular dementia; ALS, amyotrophic lateral sclerosis; FTD, frontotemporal dementia; PD, Parkinson's disease; LBD, Lewy body dementia; PMI, postmortem interval. <sup>2</sup>Other diagnoses included mild cognitive impairment (MCI; n = 3), progressive supranuclear palsy (PSP; n = 2), corticobasal degeneration (CBD; n = 1), and 1 healthy control specimen.
\*Brain weight and pH were available for 35/40 and 29/40 specimens, respectively.

## Signal-background quality assessment: GRASE and TSE acquisitions

To evaluate sequence-related signal quality, signal and background characteristics were assessed across echo times in tissue, formalin, and air regions for both GRASE32 and TSE (Figure 2). Mean tissue signal showed the expected decline with increasing TE for both sequences, while formalin signal decayed more slowly and air signal remained near zero throughout (Figure 2. A, D). With normalization to the first echo, tissue signal decay was substantially faster than formalin decay for both sequences (Figure 2. B, E); TSE additionally showed a distinct early rise in normalized air and tissue signal at the shortest echo times before the expected monotonic decay (Figure 2. E).

**Figure 2.**
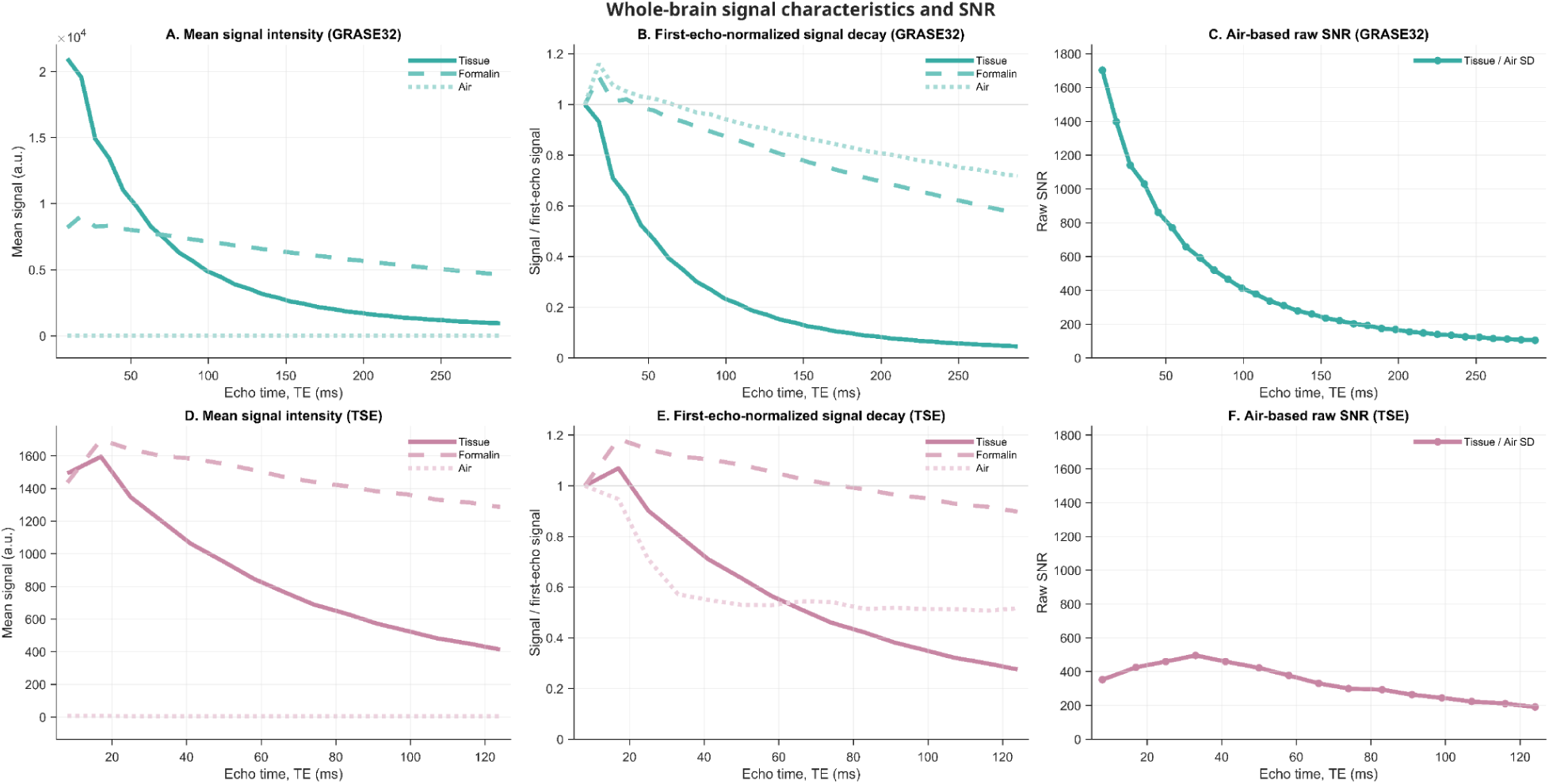
Whole-brain signal characteristics and signal-to-noise ratio for GRASE32 and TSE acquisitions. (A, D) Mean signal intensity measured in brain tissue, formalin, and air for GRASE32 and TSE, respectively. (B, E) Signal decay normalized within each subject to the corresponding first echo for tissue, formalin, and air, followed by averaging across specimens. (C, F) Raw air-based signal-to-noise ratio (SNR), calculated at each echo as the mean tissue signal divided by the standard deviation of the air signal. Curves represent group means across specimens.

## Effect of echo reduction within GRASE

Voxel-wise comparison showed very strong agreement between GRASE acquisition with the full 32-echo reconstruction (GRASE32) and the reduced 14-echo reconstruction (GRASE14) for the four derived measures (Figure 3. A). MWF showed the highest agreement between GRASE32 and GRASE14, with a Spearman correlation of ρ=0.998 (p_FDR_ < 0.001), and a mean bias of -0.07 percentage points, and an RMSE of 1.69 percentage points. IEWF similarly showed a high voxelwise correlation with ρ=0.995 (p_FDR_ < 0.001), a mean bias of -0.30 percentage points, and an RMSE of 2.60 percentage points. For the other T_2_-derived measures, T_2,M_ showed ρ=0.983 (p_FDR_ < 0.001), a mean bias of 0.52 ms, and an RMSE of 1.15 ms, whereas T_2,IE_ showed ρ=0.976 (p_FDR_ < 0.001), a mean bias of −1.51 ms, and an RMSE of 4.47 ms. The voxelwise regression slopes of GRASE32 on GRASE14 were 1.040 for MWF, 1.064 for IEWF, 1.022 for T_2,M_, and 1.039 for T_2,IE_.

**Figure 3.**
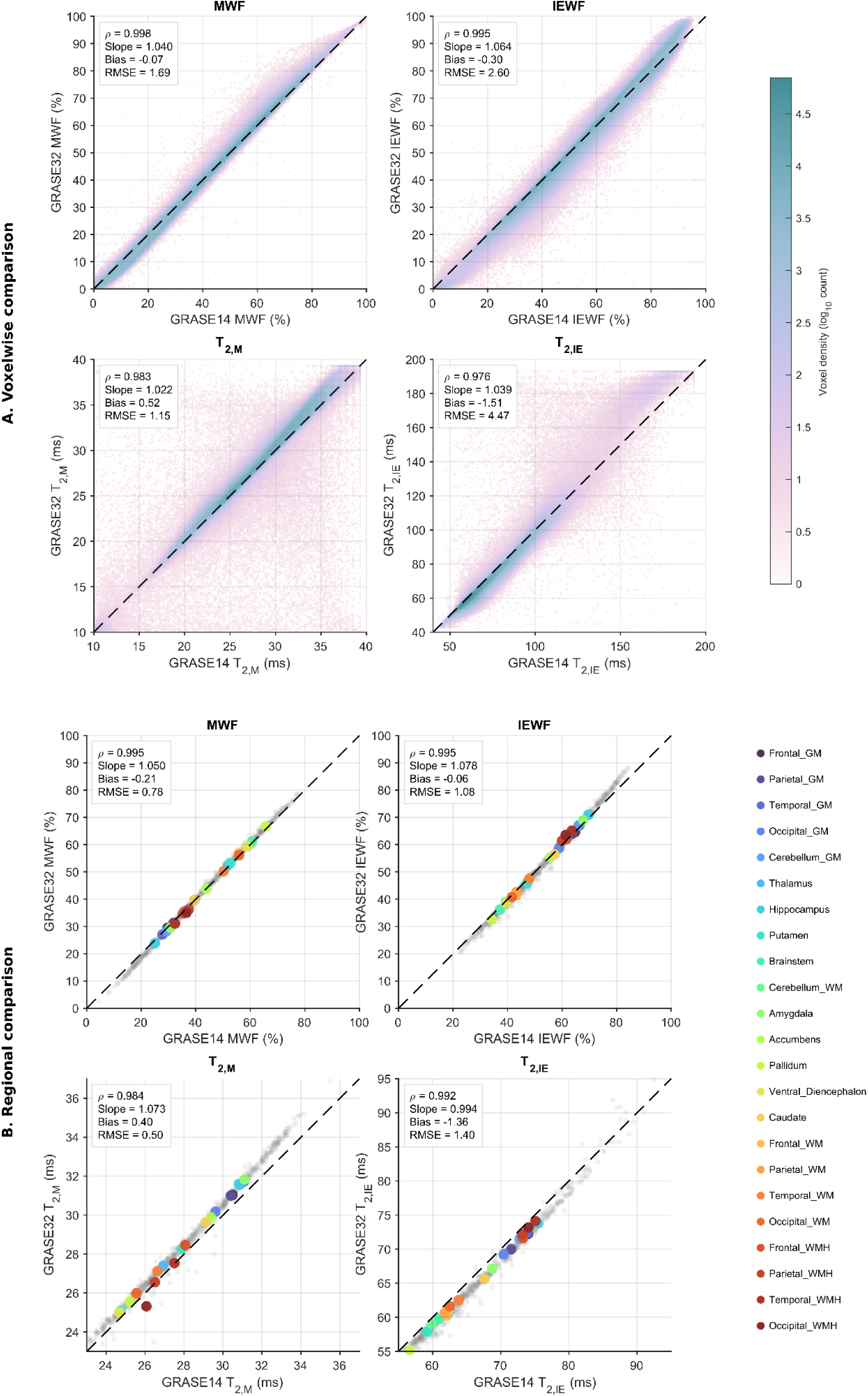
Voxelwise and regional agreement between 14-echo and 32-echo GRASE-derived quantitative maps. (A) Voxelwise comparison of GRASE14 and GRASE32 for (I) myelin water fraction (MWF), (II) intra-/extracellular water fraction (IEWF), (III) myelin-water T_2_ (T_2,M_), and (IV) intra-/extracellular water T_2_ (T_2,IE_). All valid paired voxels across specimens are shown. (B) Regional comparison of the same quantitative measures. Gray points represent individual speciment–region observations, while colored points represent the mean value for each anatomical region across specimens; colors identify regions as indicated in the legend. The dashed line denotes the line of identity. In all panels, GRASE14 is shown on the x-axis and GRASE32 on the y-axis. Insets report Pearson correlation (r), Spearman correlation (ρ), bias (GRASE32 − GRASE14), and RMSE; voxelwise panels additionally report the slope of GRASE32 regressed on GRASE14.

Regional comparisons across the 23 anatomical regions also showed high correlations between GRASE14 and GRASE32 (Figure 3. B). Spearman correlations were ρ=0.995 for MWF, ρ=0.995 for IEWF, ρ=0.984 for T_2,M_, and ρ=0.992 for T_2,IE_, with all associations remaining significant after FDR correction (p_FDR_<0.001). The regional regression slopes were 1.050 for MWF (95% CI, 1.035–1.065; p_FDR_<0.001 versus unity), 1.078 for IEWF (95% CI, 1.051–1.105; p_FDR_<0.001), 1.073 for T_2,M_ (95% CI, [1.021–1.125]; p_FDR_=0.011), and 0.994 for T_2,IE_ (95% CI, 0.970–1.019; p_FDR_=0.617). Regional mean biases were −0.21 percentage points for MWF, −0.06 percentage points for IEWF, 0.40 ms for T_2,M_, and −1.36 ms for T_2,IE_, with corresponding RMSE values of 0.78 percentage points, 1.08 percentage points, 0.50 ms, and 1.40 ms, respectively.

## Comparison of GRASE32 and TSE-derived quantitative measures

Voxelwise correspondence between GRASE32 and TSE differed across the four derived measures (Table 5; Figure 4. A). MWF showed the highest voxelwise correlation of the four measures (ρ=0.827, p_FDR_<0.001), with a mean bias of 20.50 percentage points (GRASE32 − TSE) and an RMSE of 24.60 percentage points. The voxel-density plot showed that TSE MWF values were generally lower than the corresponding GRASE32 values. IEWF showed a lower voxelwise correlation (ρ=0.593, p_FDR_<0.001), with a mean bias of −16.22 percentage points and an RMSE of 25.40 percentage points, corresponding to generally higher IEWF values with TSE than with GRASE32. Among the other T_2_-derived measures, T_2,IE_ showed a voxelwise correlation of ρ=0.768 (p_FDR_<0.001), with a mean bias of −10.24 ms and an RMSE of 17.84 ms. T_2,M_ showed the lowest voxelwise correlation (ρ=0.411, p_FDR_<0.001), with a mean bias of −3.44 ms and an RMSE of 6.81 ms. The corresponding regression slopes of GRASE32 on TSE were 1.009 for MWF, 0.575 for IEWF, 0.141 for T_2,M_, and 0.531 for T_2,IE_.

**Figure 4.**
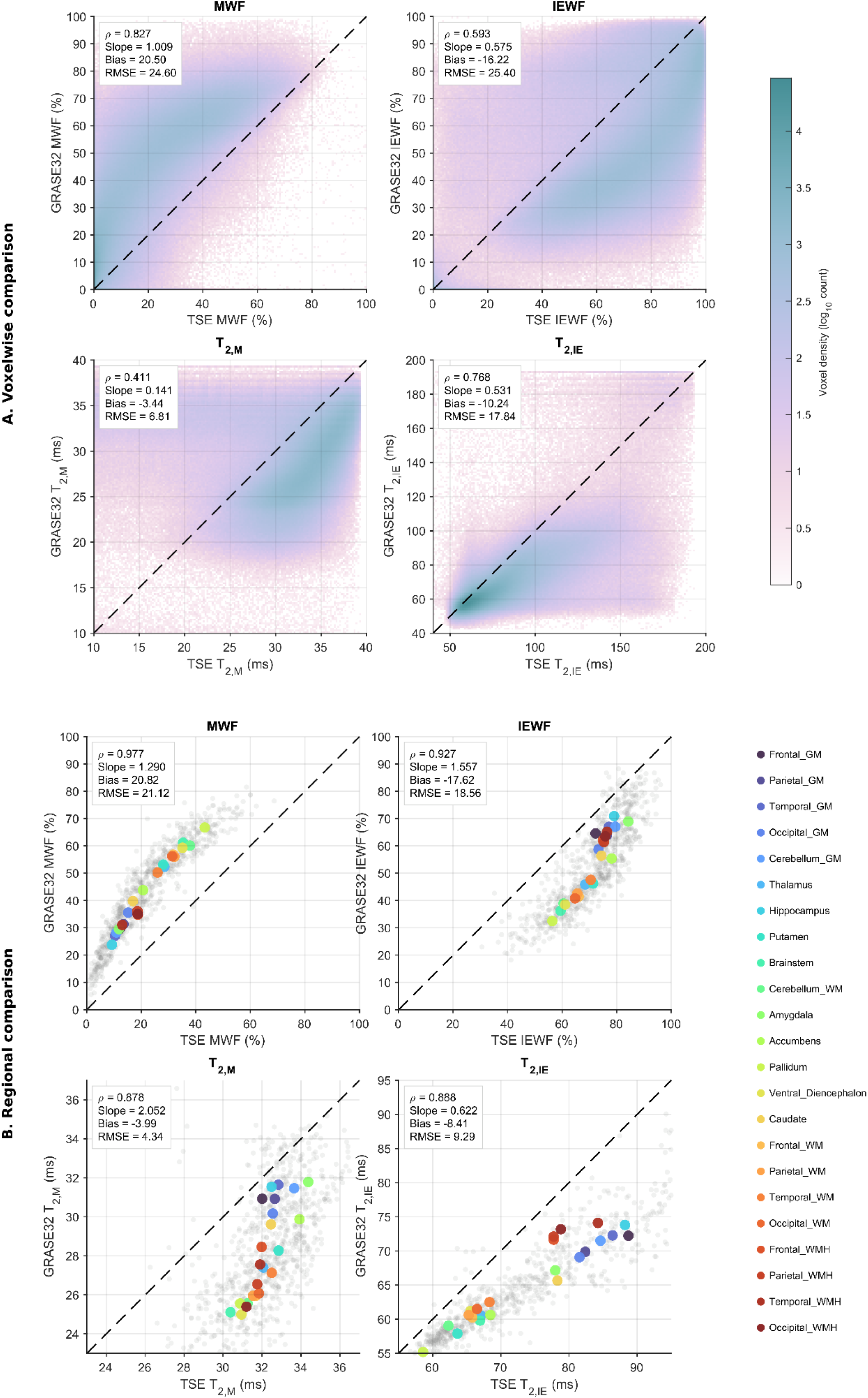
Voxelwise and regional comparison between GRASE32- and TSE-derived quantitative measures. **(A)** Pooled voxelwise distributions of MWF, IEWF, T₂,M, and T₂,IE for GRASE32 and TSE. For the voxelwise analysis, TSE quantitative maps were resampled to GRASE space, and only voxels with valid measurements in both acquisitions were included. Insets report Spearman rank correlation (*ρ*), mean bias (TSE − GRASE32), and RMSE. **(B)** Regional comparison of GRASE32 and TSE across 23 anatomical regions. Gray points represent individual specimen-region measurements, while colored points represent the mean across specimens for each anatomical region; colors identify anatomical regions consistently across measures. The dashed line indicates the line of identity. Regional correlation and error metrics were calculated across the 23 region-level mean pairs. MWF and IEWF are expressed as percentages and T₂,M and T₂,IE in milliseconds.

**Table 5.** Voxelwise and regional comparison of quantitative measures from GRASE32 and TSE.

| Statistics* | Voxelwise Analysis |  |  |  | Regional Analysis |  |  |  |
| --- | --- | --- | --- | --- | --- | --- | --- | --- |
| | MWF | IEWF | $T_{2,M}$ | $T_{2,IE}$ | MWF | IEWF | $T_{2,M}$ | $T_{2,IE}$ |
| GRASE32 | 40.99 ± 22.23 | 55.16 ± 21.04 | 28.80 ± 4.08 | 66.60 ± 15.63 | 43.37 ± 13.40 | 53.71 ± 12.24 | 28.19 ± 2.48 | 65.76 ± 6.14 |
| TSE | 20.50 ± 17.43 | 71.38 ± 20.20 | 32.24 ± 5.00 | 76.85 ± 22.21 | 22.57 ± 10.26 | 71.53 ± 7.46 | 32.83 ± 1.39 | 74.12 ± 9.34 |
| Spearman $\rho$ | 0.827 | 0.593 | 0.411 | 0.768 | 0.977 | 0.927 | 0.878 | 0.888 |
| MAE | 21.07 | 21.96 | 5.72 | 11.65 | 20.82 | 17.62 | 3.99 | 08.41 |
| RMSE | 24.60 | 25.40 | 6.81 | 17.84 | 20.12 | 18.56 | 4.34 | 9.29 |
| N of voxel/region | 2,947,692 | 2,947,692 | 2,768,796 | 2,936,504 | 23 | 23 | 23 | 23 |

\* MWF and IEWF are expressed as percentages and $T_{2,M}$ and $T_{2,IE}$ in milliseconds. For voxelwise analyses, $N$ denotes the number of paired voxels; for regional analyses, $N$ denotes the number of anatomical regions.

Regional comparisons across the 23 regions showed higher cross-sequence correlations for all four measures (Figure 4, B). Spearman correlations were ρ=0.977 for MWF, ρ=0.927 for IEWF, ρ=0.878 for T_2,M_, and ρ=0.888 for T_2,IE_, with all correlations remaining significant after FDR correction (p_FDR_<0.001). The regional regression slopes of GRASE32 on TSE were 1.290 for MWF (95% CI, 1.195–1.385; p_FDR_<0.001 versus unity), 1.557 for IEWF (95% CI, 1.291–1.823; p_FDR_<0.001), 2.052 for T_2,M_ (95% CI, 1.399–2.705; p_FDR_=0.003), and 0.622 for T_2,IE_ (95% CI, 0.527–0.717; p_FDR_<0.001). Regional mean biases were 20.82 percentage points for MWF, −17.62 percentage points for IEWF, −3.99 ms for T_2,M_, and −8.41 ms for T_2,IE_, with corresponding RMSE values of 21.12 percentage points, 18.56 percentage points, 4.34 ms, and 9.29 ms, respectively.

## TSE and GRASE associations within tissue classes

For the tissue-specific analysis, the anatomical regions were grouped into four broader tissue classes: cortical GM, deep GM, WM, and WMHs. The relationship between TSE and GRASE32 differed across tissue classes for MWF, IEWF, and T₂_,M_, but not T₂_,IE_ (Figure 5). To evaluate whether the TSE–GRASE32 relationship varied by tissue type, we fit a linear mixed-effects interaction model across the four tissue classes, allowing the TSE-to-GRASE32 slope to vary by tissue and using WM as the reference.

**Figure 5.**
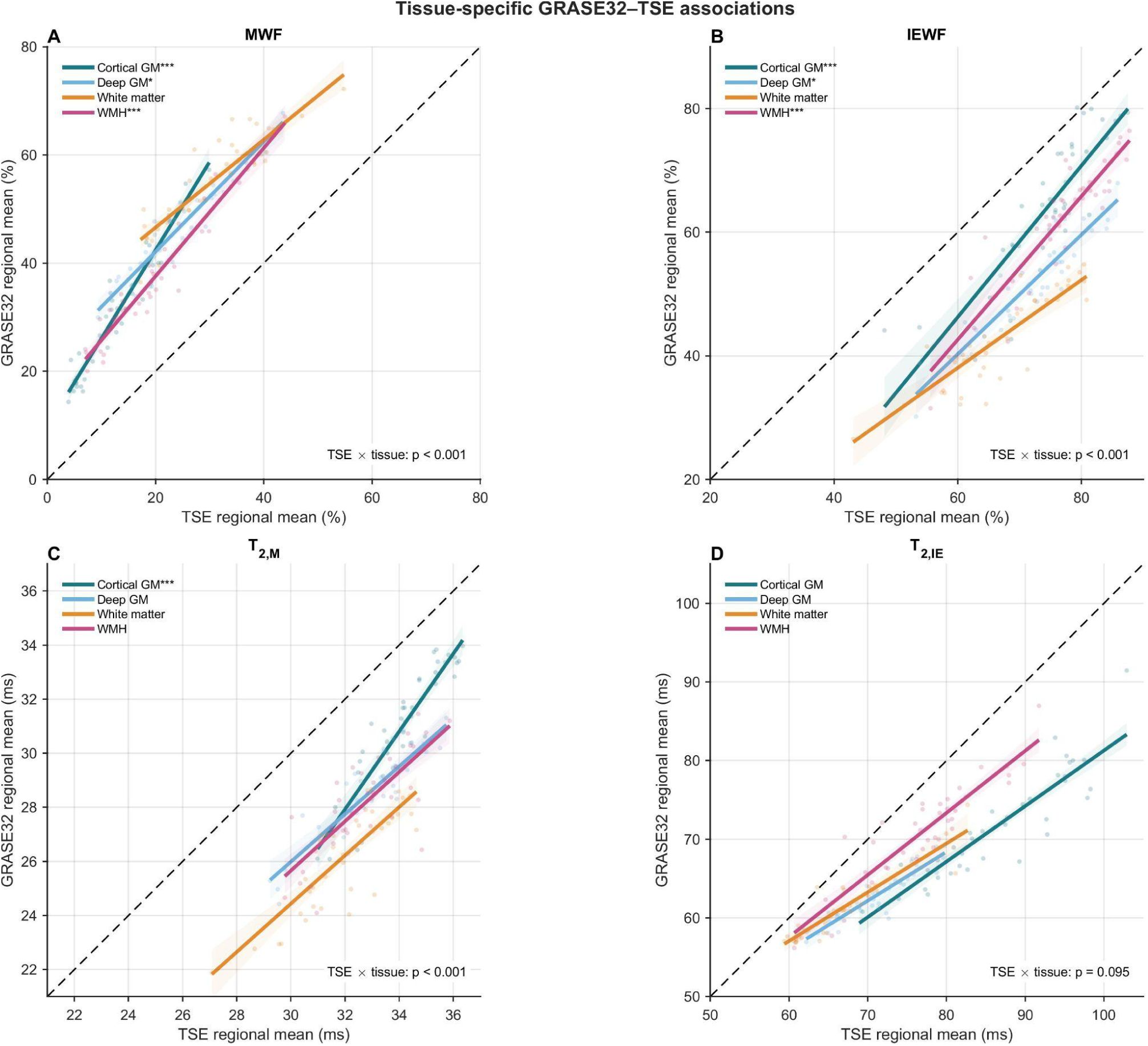
Tissue-specific GRASE32–TSE associations across quantitative MRI measures. Associations between TSE and GRASE32 measurements are shown for cortical gray matter, deep gray matter, white matter, and white matter hyperintensities (WMH) for (A) MWF, (B) IEWF, (C) T₂_,M_, and (D) T₂_,IE_. Points represent subject-level macro-region means, and colored lines show population-level fixed-effect predictions from linear mixed-effects models including TSE, tissue class, and their interaction, with subject included as a random intercept. Shaded areas indicate 95% confidence intervals, and the dashed line represents the line of identity. The TSE × tissue interaction tested whether cross-sequence slopes differed among tissue classes. Significant interactions were observed for MWF (p < 0.001), IEWF (p < 0.001), and T₂_,M_ (p < 0.001), but not T₂_,IE_ (p = 0.095). N = 40 specimens, except for WMH analyses (N = 39). *Asterisks indicate tissue-class slopes that differed significantly from the white matter reference slope after FDR correction within each measure (\**p*<0.05, \*\**p*<0.01, \*\*\**p<0.001)*.

For MWF, slopes were β = 1.63 in cortical GM, β =1.02 in deep GM, β =0.81 in WM, and β =1.19 in WMH; relative to WM, slopes were significantly higher in cortical GM (p_FDR_ < 0.001), deep GM (p = 0.016), and WMH (p_FDR_ < 0.001). For IEWF, slopes were β = 1.23, 0.97, 0.71, and 1.16; and in reference to WM, slopes were significantly higher in cortical GM (p_FDR_ < 0.001), WMH (p_FDR_ < 0.001), and deep GM (p_FDR_ = 0.027).

For T_2,M_, the cortical GM slope (β = 1.44) was significantly greater than the WM slope (β = 0.89; p_FDR_ < 0.001), whereas deep GM (β = 0.88; p_FDR_ = 0.923) and WMH (β = 0.91; p_FDR_ = 0.923) did not significantly differ from WM. For T_2,IE_, the cortical GM slope was β = 0.71, deep GM and WM slopes were β =0.62, and for WMH the slope was β = 0.79. None of the tissue-specific slopes differed significantly from WM after FDR correction (cortical GM: *p* = 0.352; deep GM: *p* = 0.989; WMH: *p* = 0.093). Consistent with these tissue-specific comparisons, the omnibus TSE × tissue interaction was significant for MWF, IEWF, and T₂,_M_ (all p_FDR_ < 0.001), but not for T₂,_IE_ (p_FDR_ = 0.095).

Correlations between TSE and GRASE32 were assessed separately within each tissue class for each of the four derived measures. TSE and GRASE32 were strongly correlated within all four tissue classes across all measures. For MWF, Spearman correlations were ρ = 0.94 in cortical GM, 0.92 in deep GM, 0.86 in WM, and 0.95 in WMH. For IEWF, correlations were ρ = 0.88, 0.90, 0.83, and 0.90 in the same four tissue classes, respectively. For T_2,M_, correlations were ρ = 0.91, 0.87, 0.81, and 0.75, and for T₂,_IE_, ρ = 0.93, 0.91, 0.89, and 0.88. All within-tissue correlations remained significant after FDR correction (p_FDR_ < 0.001).

## Impact of long-term formaldehyde exposure

Whole-brain T_2_ distributions showed systematic variation across fixation durations for both GRASE32 and TSE, with substantial overlap in the overall range of spectral profiles but sequence-dependent differences in their distribution shapes (Figure 6. A, B). In both sequences, higher whole-brain MWF was associated with longer fixation duration after adjustment for age and sex (Figure 6. C). The standardized fixation slope was 0.49 for GRASE32 and 0.39 for TSE, indicating that a one-standard-deviation increase in log-transformed fixation duration was associated with increases of 0.49 and 0.39 standard deviations in MWF, respectively. The corresponding adjusted MWF values at the mean log-fixation duration were 41.0% for GRASE32 and 19.7% for TSE. Importantly, the fixation-by-sequence interaction was significant (p_interaction_<0.001), indicating that the strength of the association between fixation duration and MWF differed between the two acquisition sequences. Specifically, the fixation-related increase in MWF was greater for GRASE32 than for TSE.

**Figure 6.**
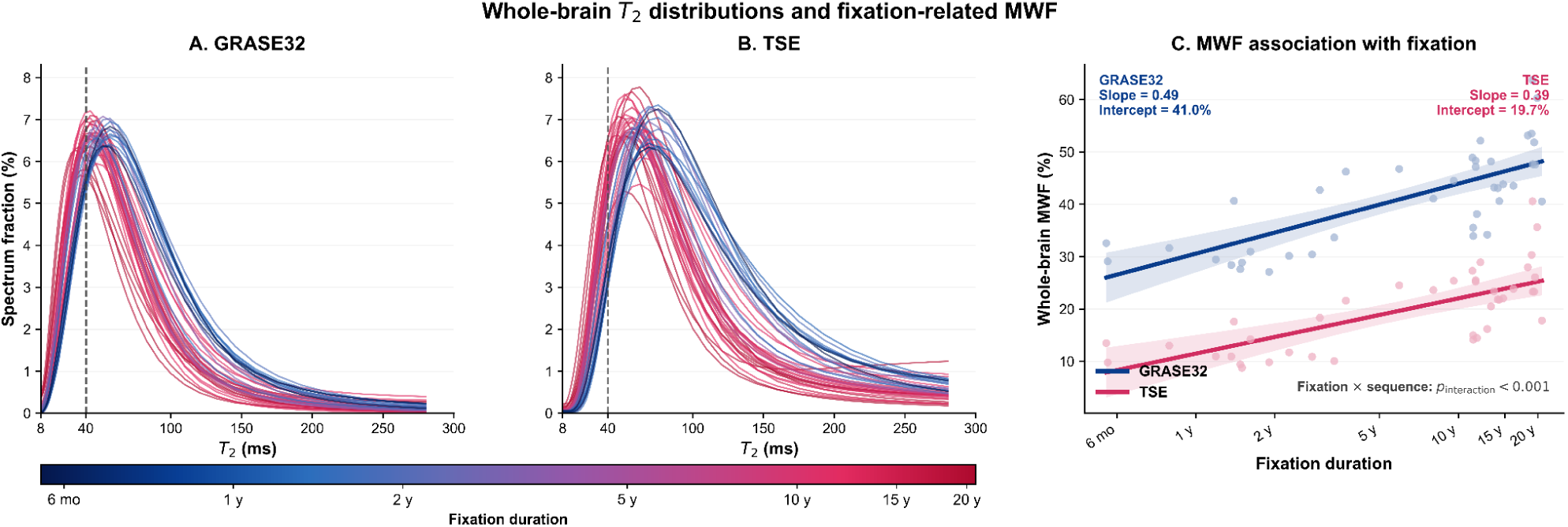
Whole-brain T_2_ distributions and fixation-related MWF in GRASE32 and TSE. Whole-brain multicomponent T_2_ distributions are shown for (A) GRASE32 and (B) TSE, with individual specimen spectra color-coded by fixation duration. Spectra are displayed over 8–300 ms, and the dashed vertical line indicates the 40-ms threshold used to define the short-T_2_ component contributing to the myelin water fraction (MWF). (C) Association between fixation duration and whole-brain MWF for GRASE32 and TSE. Fixation duration was log-transformed and standardized, and MWF was standardized using a common pooled scale across both sequences. Solid lines show age- and sex-adjusted model fits and shaded regions indicate 95% confidence intervals, back-transformed to the original MWF percentage scale. Reported slopes are standardized fixation effects, and intercepts correspond to adjusted predicted MWF at the mean log-transformed fixation duration. The fixation-by-sequence interaction p-value tests whether the fixation–MWF association differs between GRASE32 and TSE.

## MWF Prediction of GRASE32 from TSE

Six models were trained to predict GRASE32 MWF from the TSE data. Prediction performance improved as model flexibility and TSE-derived information increased (Table 6). The simple linear model showed the lowest overall performance, with a median RMSE of 12.58, CCC of 0.779, and masked SSIM of 0.607. Allowing a nonlinear relationship between TSE and GRASE32 MWF through a quadratic term improved performance, reducing the median RMSE to 11.46 and increasing the CCC and masked SSIM to 0.822 and 0.656, respectively. The random forest model using the same primary TSE MWF predictor (Model 1) provided a further modest improvement, with a median RMSE of 11.33, CCC of 0.827, and masked SSIM of 0.676.

**Table 6.** Cross-validated performance of TSE-to-GRASE32 MWF prediction models.

| Model | Predictors | RMSE | MAE | CCC | Pearson<br>r | SD<br>ratio | Bias | Masked<br>SSIM | Median<br>improvement<br>vs null (%) |
| --- | --- | --- | --- | --- | --- | --- | --- | --- | --- |
| Linear | TSE_MWF, regions | 12.58 | 10.33 | 0.779 | 0.829 | 0.856 | -0.48 | 0.607 | 39.98 |
| Quadratic | TSE_MWF, TSE_MWF<sup>2</sup>, regions | 11.46 | 9.05 | 0.822 | 0.844 | 0.845 | 0.33 | 0.656 | 43.56 |
| Model 1 | MWF, regions | 11.33 | 8.69 | 0.827 | 0.848 | 0.862 | 0.56 | 0.676 | 44.59 |
| Model 2 | MWF, IEWF, T2M, T2IE, regions | 9.02 | 7.09 | 0.893 | 0.909 | **0.896** | 1.29 | 0.793 | 56.08 |
| Model 3 | 12 spectrum PCs, regions | 8.88 | 7.10 | **0.896** | **0.911** | 0.892 | 1.39 | 0.799 | **56.56** |
| Model 4 | 12 spectrum PCs, MWF, T2M, regions | **8.87** | **7.07** | **0.896** | **0.911** | 0.893 | 1.42 | **0.802** | 56.55 |

Values represent median specimen-level performance across the held-out specimens. Predictions were generated using 10-fold specimen-level cross-validation. RMSE, root mean squared error; MAE, mean absolute error; CCC, concordance correlation coefficient; SD ratio, ratio of predicted to observed GRASE32 MWF standard deviation; SSIM, structural similarity index measure. Improvement relative to the null model was calculated from specimen-level RMSE.

A substantially larger improvement was observed when additional TSE-derived information was incorporated. Model 2, which added IEWF, T_2,M_, and T_2,IE_ to MWF and regional anatomy, reduced the median RMSE to 9.02 and the MAE to 7.09, with a CCC of 0.893, Pearson r=0.909, an SD ratio of 0.896, a bias of 1.29 percentage points, and a masked SSIM of 0.793. This corresponded to a 56.08% improvement over the null model. Model 3 replaced the derived T_2_ measures with 12 principal components from the normalized TSE T_2_ distribution and achieved a slightly lower median RMSE of 8.88. Its MAE was 7.10, with a CCC of 0.896, Pearson r=0.911, an SD ratio of 0.892, a bias of 1.39 percentage points, and a masked SSIM of 0.799; the median improvement over the null model was 56.56%. Model 4 combined the 12 spectral principal components with TSE MWF, T_2,M_, and regional anatomy. It had the lowest median RMSE of 8.87 and MAE of 7.07, together with a CCC of 0.896, Pearson r=0.911, an SD ratio of 0.893, a bias of 1.42 percentage points, and a masked SSIM of 0.802. The corresponding improvement over the null model was 56.55%. Model 4 was nonetheless selected as the final cross-sequence prediction model based on its marginally best overall performance (Table 6, Figure 8). Relative to the specimen-specific null model, the median RMSE improvement was 44.59% for Model 1 and 56.08–56.56% for Models 2–4. For each of Models 1–4, RMSE was lower than the corresponding null-model RMSE in 38 of 40 held-out specimens.

**Figure 7.**
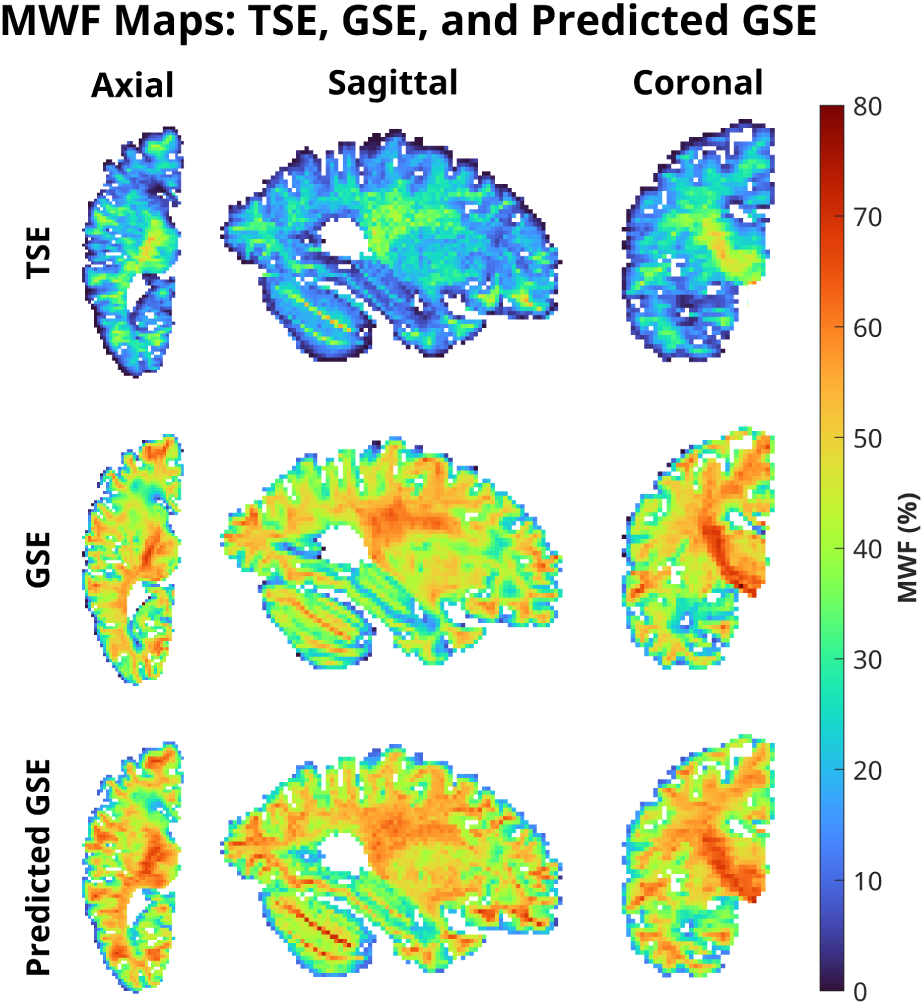
Representative MWF maps from TSE, observed GRASE32, and predicted GRASE32. Axial, sagittal, and coronal views are shown for a representative held-out specimen. TSE-derived MWF maps are shown alongside the corresponding observed GRASE32 MWF maps and out-of-fold GRASE32 MWF predictions from the final selected model. All maps are displayed using the same spatial support and color scale (0–80% MWF).

**Figure 8.**
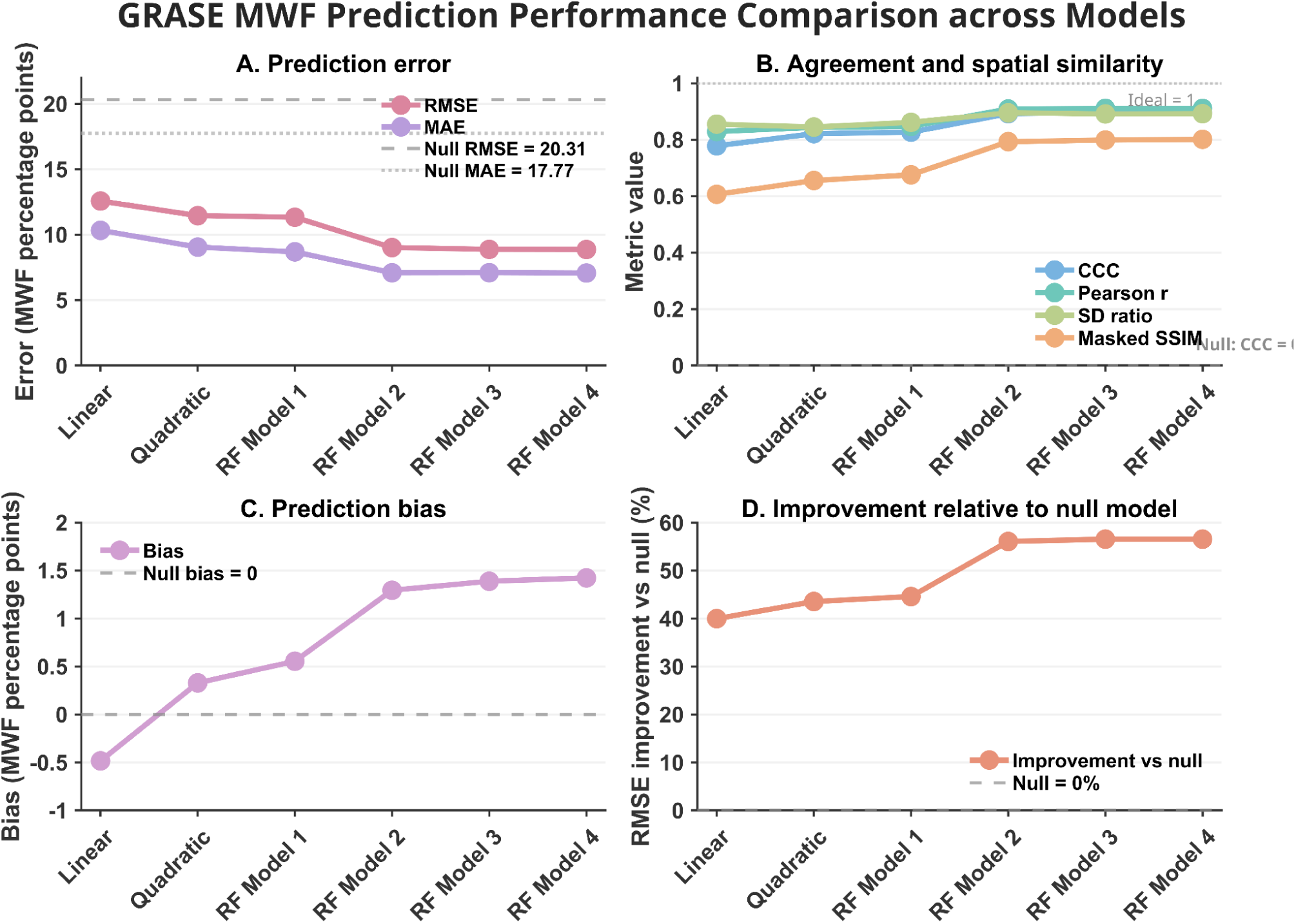
Comparison of GRASE32 MWF prediction performance across model configurations. Model performance was evaluated using held-out predictions from 10-fold specimen-level cross-validation. Values represent median specimen-level metrics across the 40 specimens. **(A)** Prediction error, quantified by root mean squared error (RMSE) and mean absolute error (MAE). **(B)** Agreement and spatial similarity, assessed using the concordance correlation coefficient (CCC), Pearson correlation coefficient (r), predicted-to-observed standard deviation (SD) ratio, and masked structural similarity index measure (SSIM); the dashed line at 1 indicates the ideal SD ratio. **(C)** Mean prediction bias, with zero indicating no systematic over- or underestimation. **(D)** Percentage improvement in RMSE relative to the specimen-specific null model.

Predicted GRASE32 MWF was compared with observed GRASE32 MWF at both voxelwise and regional levels (Figure 9). At the voxelwise level, the Spearman correlation between predicted and observed MWF was ρ=0.877. The corresponding regression had a slope of 1.009, with a mean bias of 0.03 percentage points (predicted minus observed) and an RMSE of 10.58 percentage points. At the regional level, the Spearman correlation calculated across the 23 regional mean pairs was ρ=0.978 (p<0.001). The regression observed on predicted regional MWF had a slope of 1.075 (95% CI, [0.994,1.156]), which did not differ significantly from unity (p=0.069). The regional mean bias was 0.59 percentage points, and the RMSE was 2.40 percentage points.

**Figure 9.**
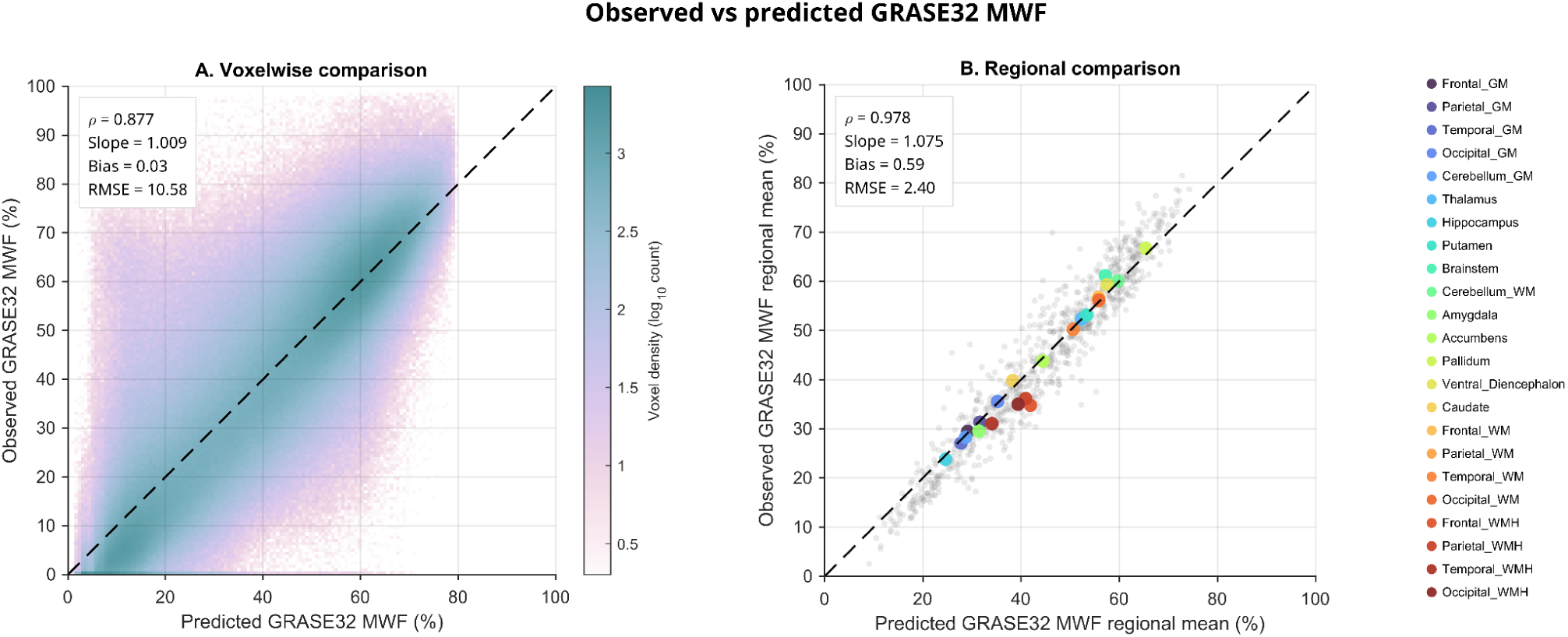
Voxelwise and regional correspondence between observed and predicted GRASE32 MWF. **(A)** Voxelwise comparison of observed GRASE32 MWF and out-of-fold predictions from the selected random forest model across 40 specimens. Color indicates the log-transformed number of voxel pairs within each bin. **(B)** Regional comparison across 23 anatomical regions. Gray points represent subject-specific regional means, and colored points represent the mean across subjects for each region. Dashed lines indicate the line of identity. Spearman’s ρ\rho, regression slope, mean bias (predicted − observed), and RMSE are shown in each panel.

## Discussion

The present study evaluated two distinct MR sequences, 3D GRASE and two-dimensional multi-echo TSE, for multicomponent T_2_ relaxometry of formaldehyde-fixed postmortem human brain specimens, with a focus on MWF, IEWF, and their corresponding T_2_ components (T_2,M_ and T_2,IE_). Both sequences provided robust multicomponent T_2_ characterization across the imaged hemispheres, with the four derived measures showing anatomical patterns consistent with expected differences across tissue types. We assessed SNR across the echo train of each sequence as an acquisition-level quality-control measure, and verified that mean tissue signal remained above the noise floor for reliable multicomponent T_2_ analysis. Comparisons between GRASE- and TSE T_2_-derived measures at the voxelwise, regional, and tissue-class levels showed moderate to strong associations, and echo reduction comparisons between the full 32-echo GRASE reconstruction and a truncated 14-echo reconstruction, matched to the TSE echo-time range, showed minimal impact of echo truncation. Finally, GRASE32 MWF could be predicted with high accuracy based on multicomponent T_2_ information derived from the TSE sequence.

Adequate SNR is particularly important for multicomponent T_2_ analysis, as loss of signal at later echoes can reduce the stability of the recovered T_2_ distribution and bias compartment estimates (30). In our study, the use of a 2D TSE sequence with higher spatial resolution and fewer echoes made it particularly important to verify that the later echoes were not approaching the noise floor. Because GRASE and TSE have intrinsic acquisition differences, their absolute SNR values cannot be directly compared (31). We therefore evaluated signal behavior independently across the echo train of each sequence as an acquisition-level quality-control measure. TSE showed the expected signal decay with increasing echo time, and the mean signal across specimens remained above the noise floor throughout the sampled echo range. We also observed an early-echo signal increase in TSE, consistent with imperfect refocusing characteristics and/or stimulated-echo effects (7), rather than a true SNR gain; this pattern was not present in GRASE32. Measurements from ROIs placed in the surrounding formalin showed a slower signal decay, consistent with a longer apparent T_2_ relative to fixed tissue (32). Overall, this analysis provided internal validation of the signal quality supporting the multicomponent T_2_ estimates used in the subsequent cross-sequence comparisons and prediction analyses.

Reducing the GRASE echo train from 32 to 14 echoes had a small effect on the multicomponent T_2_ derived measures. Truncating the acquisition from 32 to 14 echoes produced very high voxelwise and regional correspondence across all four derived measures. The water-fraction measures, particularly MWF, were especially stable, with only minimal bias between the two reconstructions. T_2,M_ and T_2,IE_ also remained strongly correlated, although these two measures showed slightly greater sensitivity to echo truncation than the fraction estimates. Small systematic scaling differences were present, with regression slopes slightly above unity for most measures, indicating that echo reduction was not entirely neutral despite the overall high agreement. At the regional level, correspondence was similarly strong, with negligible differences in the water-fraction measures and similarly small directional shifts in the mean T_2_ estimates (T_2,M_, T_2,IE_). These findings are notable because multicomponent T_2_ imaging has traditionally relied on relatively long multi-echo acquisitions to characterize the complex relaxation behavior of brain tissue (1,11,29). More recent studies, however, have shown that shorter acquisitions can still provide reliable MWF estimates that agree well with standard approaches, including dual-echo spin-echo methods (34), approximately 12 nonlinear echo times (35), and protocols reduced to as few as 6 echoes (36). Our findings extend this observation to formaldehyde-fixed postmortem tissue and suggest that the complete 32-echo range may not be necessary to preserve the main multicomponent T_2_ measures examined here. The choice of echo-time range and spacing should therefore be tailored to the intended measurement and reconstruction approach (13). Although GRASE has demonstrated high reproducibility in vivo (37), postmortem tissue may particularly benefit from shorter initial TEs and reduced echo spacing because fixation shortens tissue T_2_. In this setting, acquisition time devoted to very late echoes could be redirected toward improving other acquisition parameters. Longer echo times could still be valuable when sensitivity to the long-T_2_ free or quasi-free water compartment (>200 ms), mainly associated with increased extracellular or relatively mobile water (5), is required. However, characterization of this compartment was outside the primary scope of the present protocol.

The comparison between TSE and GRASE32 showed that the two sequences captured related but quantitatively different multicomponent T_2_ information. MWF showed the strongest voxelwise correspondence, yet TSE consistently produced lower MWF values than GRASE32, indicating a systematic sequence-dependent difference in the estimated short-T_2_ fraction. This reduction in TSE-derived MWF was accompanied by higher IEWF values. Because MWF and IEWF represent complementary fractional contributions of the same recovered T_2_ distribution, these shifts are inherently linked: a smaller proportion of signal assigned to the short-T_2_ pool will increase the proportion assigned to the intermediate-T_2_ compartment. The reciprocal MWF–IEWF pattern therefore likely reflects differences in compartment separation between TSE and GRASE32 rather than opposing biological effects. This interpretation is further supported by the less distinct short-T_2_ component observed in the TSE spectra, suggesting that differences in temporal sampling and sequence-specific signal formation (7) may influence how signal is partitioned between adjacent T_2_ compartments. Similar cross-sequence correspondence in MWF has also been reported previously. Piredda et al. (9) found a correlation of r=0.83 with a small mean bias between GRASE and conventional MESE-derived MWF, supporting the observation that different T_2_-based acquisition schemes can produce similar overall MWF patterns despite differences in acquisition methodology.

The corresponding T_2_ measures (T_2,M_ and T_2,IE_) also showed different sensitivity to acquisition sequence. T_2,IE_ maintained relatively strong voxelwise correspondence between TSE and GRASE32 despite a systematic shift toward higher TSE-derived values, whereas T_2,M_ showed the weakest voxelwise correspondence despite relatively small absolute differences between the sequences. T_2,M_ represents the geometric mean relaxation time of the short-T_2_ myelin-water component and therefore depends directly on reliable separation of this component from the intermediate-T_2_ pool (13,21,38) . Its weaker cross-sequence correspondence may reflect greater sensitivity of the short-T_2_ estimate to sequence-related differences. In contrast, T_2,IE_ showed stronger correspondence, suggesting greater stability of the intermediate-T_2_ component across the acquisitions despite differences in absolute values. This relative stability is consistent with previous reports of high scan–rescan and intersite reproducibility of T_2,IE_, which has also been proposed as a quantitative marker of changes in the tissue-water environment and water mobility (21,39).

Regional correlations showed substantially higher TSE–GRASE32 correspondence across all four T_2_-derived measures, with particularly large improvements for T_2,M_ and IEWF, which showed weaker correspondence at the voxel level. MWF also showed very high regional correspondence, while T_2,IE_ remained strongly associated across regions. This suggests that a substantial component of the disagreement between acquisitions occurs at the local voxel scale and that TSE preserves the relative anatomical variation captured by GRASE32 more reliably than it reproduces individual voxel values. Regional aggregation likely reduces sensitivity to local fitting instability, noise, partial-volume effects, and residual spatial mismatch between acquisitions. However, the stronger regional correspondence should not be interpreted as quantitative equivalence between TSE and GRASE32, as systematic differences in the regional means remained and the regional regression slopes differed significantly from unity for all four measures. Overall, the relationship between TSE- and GRASE32-derived measures was more stable at the regional scale than at the voxel level.

When the anatomical regions were grouped into four broader tissue classes (cortical GM, deep GM, WM, and WMHs) the TSE–GRASE32 relationship differed significantly across tissue types for MWF, IEWF, and T_2,M_, but not for T_2,IE_. For MWF and IEWF, the cross-sequence relationship in cortical GM, deep GM, and WMH differed from that in WM, whereas for T_2,M_ the difference was driven by cortical GM alone. For T_2,IE_, the relationship was comparatively consistent across tissue classes. These differences occurred despite strong within-tissue correlations for all four measures, indicating that the two sequences track variation within individual tissue classes well, but that their relationship is not adequately described by a single global transformation across all tissues.

The corresponding changes in MWF and IEWF are consistent with their complementary fractional nature. Both are derived from the same recovered T_2_ distribution, such that sequence-related differences in assignment of signal to the short-T_2_ component can affect both fractions simultaneously. The stronger tissue dependence of the fractional measures, together with the more limited effect for T_2,M_ and the absence of a significant effect for T_2,IE_, suggests that sequence-related differences may arise more strongly from the separation and partitioning of signal between T_2_ components than from estimation of the relaxation time of the intermediate-T_2_ pool.

These findings also have practical implications for cross-sequence harmonization. The tissue-dependent relationship observed for MWF suggests that a calibration derived from one tissue type will not necessarily generalize directly to another. Studies pooling measurements across tissue classes, or comparing tissue classes using different acquisition sequences, may therefore need to account for tissue type during calibration to avoid introducing systematic sequence-related differences. A similar consideration applies to IEWF, whereas T_2,IE_ showed no significant evidence of tissue-dependent cross-sequence differences.

Our findings in positive association between fixation duration and MWF is consistent with previous longitudinal observations during early formalin fixation. Shatil et al. (40) demonstrated progressive increases in MWF accompanied by reductions in T_1_ and T_2_ during the first ∼43 days of fixation, with changes persisting through the latest measurement. We extend this observation to specimens exposed to formaldehyde for substantially longer periods, suggesting that fixation-related differences in multicomponent T_2_ measures remain detectable beyond the early fixation interval. Such increases should not be interpreted as increased myelin content. Since aldehyde fixation substantially modifies tissue-water relaxation and exchange, it can consequently alter the relative signal assigned to different portions of the T_2_ spectrum (41). Both TSE and GRASE32 captured the same direction of the fixation effect on this “apparent MWF” but fixation-by-sequence interaction showed that its magnitude was sequence-dependent, with a stronger increase measured by GRASE32. This finding is consistent with the different recovery of the short-T_2_ component by the two acquisitions, showing a greater fixation sensitivity observed with GRASE32, which may reflect both improved sensitivity to the short-T_2_ component and greater sensitivity to fixation-induced redistribution of the T_2_ spectrum. Histology studies showed a correlation between GRASE and myelin coloration with mean R^2^ = 0.63 (33) therefore, reproducible studies with both methods still needed (42).

Despite the systematic and tissue-dependent differences between TSE and GRASE32, correlations between the two sequences were consistently high within tissue classes and across anatomical regions, indicating that both sequences capture substantial overlapping tissue-related information. This motivated us to examine whether GRASE32 MWF could be predicted directly from the multicomponent T₂ information available in the 15-echo TSE acquisition. We first tested whether the cross-sequence mapping could be described parametrically, beginning with a linear model and then adding a quadratic term, which better reflected the curvature apparent in the observed TSE–GRASE32 relationship. The quadratic model improved on the linear model, but both performed substantially worse than models incorporating additional TSE-derived information. We therefore used random forests for the subsequent models, allowing nonlinear relationships and interactions among predictors to be captured without specifying their functional form in advance. Given the tissue-dependent TSE–GRASE32 relationships observed in the preceding analyses, the 23 anatomical regions were included as categorical predictors throughout the prediction framework.

Prediction improved when the additional multicomponent T₂ measures, IEWF, T₂,_M_, and T₂,_IE_, were included alongside MWF for TSE, and improved slightly further when these derived measures were replaced by principal components of the full normalized TSE T₂ distribution. Combining the spectral components with TSE MWF and T_2,M_ gave the best overall performance, although the difference from the spectrum-only model was marginal. The similar high performance of the spectral representation and the explicitly derived measures suggests that the information required to recover GRASE32 MWF is present in the TSE T_2_ distribution itself, and that the summary measures almost completely capture the relevant information contained within it (R^2^ of 0.807 compared to 0.811). Their better performance than TSE MWF alone further indicates that information useful for predicting GRASE32 MWF is distributed across the T_2_ spectrum and is not fully captured by the short-T_2_ fraction alone. Together with the systematic MWF and IEWF differences observed between sequences, this supports the interpretation that at least part of the cross-sequence discrepancy reflects differences in how the recovered T₂ signal is separated and partitioned into components, rather than an absence of myelin-related information in the TSE acquisition.

Several limitations should be considered. First, the study was performed in formaldehyde-fixed postmortem tissue, where relaxation properties differ substantially from those in vivo, and therefore, the quantitative relationships observed between TSE and GRASE32 should not be assumed to transfer directly to in vivo acquisitions. Second, TSE and GRASE differed in several acquisition characteristics, including dimensionality, spatial resolution, echo sampling, and signal formation. The present comparison therefore characterizes the combined sequence-level differences rather than isolating the contribution of any single acquisition parameter. Although images were spatially aligned for voxelwise analyses, residual registration and resampling effects may also contribute to local disagreement between sequences. Furthermore, tissue-dependent relationships may also be impacted by post-mortem fixation effects. Formaldehyde penetrates brain tissue progressively from superficial toward deeper regions, producing spatially and temporally dependent changes in T_2_ relaxation and MWF (32,40). Consequently, regional differences in the degree of fixation may contribute additional variability to multicomponent T_2_ measures in post-mortem specimens, although our data cannot separate those two effects. Finally, the prediction models were evaluated using specimen-wise cross-validation within the same postmortem cohort and require validation in an independent dataset. Importantly, the prediction target was GRASE32-derived MWF rather than an independent histological measure of myelin; the models therefore demonstrate cross-sequence mapping rather than biological validation of either measurement.

In summary, TSE and GRASE32 captured strongly related multicomponent T₂ information in fixed postmortem human brain tissue, but with systematic and tissue-dependent quantitative differences across derived measures. Reducing the GRASE echo train from 32 to 14 echoes had only a small effect on MWF, IEWF, T₂_,M_, and T₂_,IE_, indicating that substantially shorter echo sampling can preserve the principal multicomponent estimates under these acquisition conditions. We then evaluated a 15-echo TSE acquisition with a comparable T₂-sampling range and found that, despite systematic differences from GRASE32, it retained substantial overlapping information, as reflected by strong regional correspondence and the successful prediction of GRASE32 MWF from both TSE-derived measures and the full T₂ distribution. Together, these findings suggest that the full 32-echo GRASE acquisition may not be necessary to obtain equivalent multicomponent T₂ measures in fixed postmortem tissue, and that a 15-echo TSE acquisition with a comparable echo range can capture much of the information relevant to GRASE32-derived MWF. However, the systematic and tissue-dependent differences between sequences indicate that direct interchangeability should not be assumed and that cross-sequence mapping or calibration is required when quantitative equivalence is desired.

## Acknowledgements

We would like to acknowledge Dr. Gerald R. Moran (Head of Research, Innovation and Scientific Engagement, Siemens Healthcare Limited) who provided the GRASE sequence through a research application. We would also like to thank Danae Lussier-Dumouchel and Elena Drobotea for their help in MRI data acquisition. This project was supported by research funds from the Healthy Brains for Healthy Lives (HBHL), the Quebec BioImaging Network (QBIN), the Natural Sciences and Engineering Research Council of Canada (NSERC), the Canadian Institutes of Health Research (CIHR), Brain Canada, and ALS Canada-Brain Canada.

MD reports receiving research funding from the CIHR (191303, 198104, and 213169), NSERC discovery grant (RGPIN-2023-04038), FRQS (https://doi.org/10.69777/330750), Alzheimer’s Society Research Program (ASRP), Brain Canada, and Tier-2 Canada Research Chair in Vascular and Neurodegenerative Disorders of Aging. YZ reports receiving research funding from the FRQS (https://doi.org/10.69777/379799), NSERC discovery grant (RGPIN-2023-04218), ALS-Canada Brain Canada, and CIHR (195671). The Douglas Brain Bank is supported by a platform support grant from Brain Canada. RM and WAG are supported by scholarships from the FRQS. ZA is supported by a doctoral NSERC Vanier scholarship.

